# Sequestration of RBM10 in Nuclear Bodies: Targeting Sequences and Biological Significance

**DOI:** 10.1101/516831

**Authors:** Ling-Yu Wang, Sheng-Jun Xiao, Hiroyuki Kunimoto, Kazuaki Tokunaga, Hirotada Kojima, Masatsugu Kimura, Takahiro Yamamoto, Naoki Yamamoto, Zhao Hong, Koji Nishio, Tokio Tani, Koichi Nakajima, Kishiko Sunami, Akira Inoue

**Author notes:** Corresponding author: Akira Inoue, Department of Otolaryngology, Osaka City University Graduate School of Medicine, 1-4-3 Asahi-machi, Abeno-ku, Osaka 545-8585, Japan. These authors contributed equally to this work (Ling-Yu Wang, Sheng-Jun Xiao).

## Abstract

RBM10 is an RNA-binding protein that regulates alternative splicing (AS). It localizes to the extra-nucleolar nucleoplasm and S1-1 nuclear bodies (NBs) in the nucleus. We investigated the biological significance of this localization in relation to its molecular function. Our analyses, employing deletion mutants, revealed that RBM10 possesses two S1-1 NB-targeting sequences (NBTSs), one in the KEKE motif region and another in the C_2_H_2_ Zn finger (ZnF). These NBTSs act synergistically to localize RBM10 to S1-1 NBs. The C_2_H_2_ ZnF not only acts as an NBTS, but is also essential for AS regulation by RBM10. Moreover, RBM10 does not participate in S1-1 NB formation, and without alterations of RBM10 protein levels, its NB-localization changes, increasing as cellular transcriptional activity declines, and vice versa. These results indicate that RBM10 is a transient component of S1-1 NBs and is sequestered in NBs via its NBTSs when cellular transcription decreases. We propose that the C_2_H_2_ ZnF exerts its NB-targeting activity when RBM10 is unbound by pre-mRNAs, and that NB-localization of RBM10 is a mechanism to control its AS activity in the nucleus.

Note that the previous title of this manuscript was Targeting of RBM10 to S1-1 Nuclear Bodies: Targeting Sequences and its Biological Significance.

## Introduction

RBM10 (RNA-binding motif protein 10, previously called S1-1) is an RNA-binding protein [1]. It regulates alternative splicing (AS) of various pre-mRNAs [2] via exon skipping, as shown for the *Drosophila* protein Discs large homolog 4 [3], the Notch pathway regulator NUMB [2, 4], apoptosis-related protein Fas [5], and so on. Loss of RBM10 function causes TARP syndrome (Talipes equinovarus, Atrial septal defect, Robin sequence, and Persistent left superior vena cava), which is characterized by various developmental anomalies, such as cleft palate and malformation of the heart [6, 7], as well as other diseases, including lung adenocarcinoma [8, 9] (reviewed in [10] and [11]).

Nuclear bodies (NBs) are membrane-less compartments formed by molecular condensation in the nucleus. There are various types of NBs, and their formation, compositions, and roles in nuclear functions have been studied (reviewed in [12] and [13]). While some types of NBs, such as Cajal bodies, nuclear speckles, and paraspeckles, have been thoroughly investigated, the biological significance of S1-1 NBs and their relationship to RBM10 remain largely unknown. RBM10 localizes to nuclear domains called S1-1 NBs and S1-1 granules. S1-1 NBs are ~0.5–2.0 μm in size, and 10–30 NBs are generally present per nucleus. S1-1 granules are much smaller and much more abundant, and likely localize to the peri-chromatin fibrils that comprise transcription and splicing sites [14]. These S1-1 nuclear domains undergo dynamic changes in response to transcriptional states of cells. When cellular transcription declines globally, S1-1 granules diminish and S1-1 NBs enlarge accumulating RBM10 [14].

To better understand RBM10 localization to S1-1 NBs, we identified the S1-1 NB-targeting sequences (NBTSs) in RBM10 that participate in its targeting and localization to S1-1 NBs, and explored the significance of this localization in relation to its AS regulatory function.

## Results

### Sequences responsible for targeting of RBM10 to S1-1 NBs

Wild-type and deletion mutants of an 852-amino acid (aa) isoform of RBM10 (Figure 1A), linked in-frame to the N-terminus of EGFP, were expressed in ARL (adult rat liver epithelial) cells, and their subcellular localization was examined with EGFP fluorescence and by staining with an anti-S1-1/RBM10 antiserum. The antibody, which detects endogenous RBM10, recognizes an epitope in aa 8–63 of RBM10 [14] (Supplementary Figure S1). In contrast to the aa 4–414 deletion, the aa 4–742 deletion completely abolished localization of RBM10 to S1-1 NBs, suggesting that an NBTS is present in aa 415–742 (Figure 1B).

**Figure 1.**
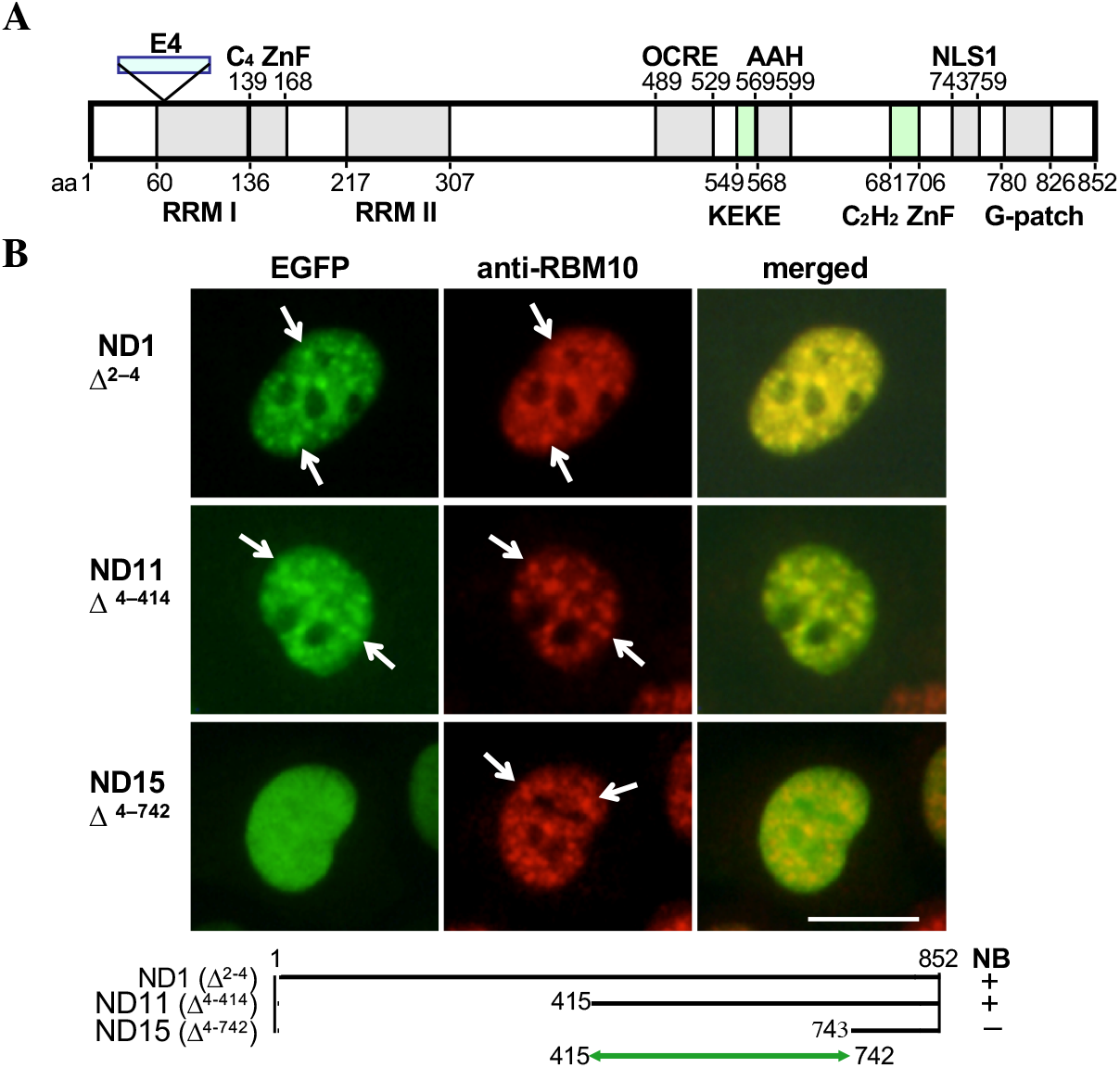
RBM10 targets NBs. **A.** Structure of the 852-aa isoform of RBM10. RRMs I and II: <u>R</u>NA <u>r</u>ecognition <u>m</u>otifs I and II, respectively; E4: a 78-aa region encoded by exon 4 and expressed in the 930-aa isoform; C_4_ ZnF and C_2_H_2_ ZnF: Cys_4_-type and Cys_2_His_2_-type ZnFs, respectively; NLS1: the principal nuclear localization sequence [16]; G-patch: a motif sometimes found in eukaryotic RNA-processing proteins [28, 29]. Other motifs, OCRE, KEKE motif, and amphipathic α-helix (AHH), are described in Figure 3. **B.** Presence of an NBTS in aa 415–742. Plasmids ND1, ND11, and ND15 were derived from pRBM10/EGFP-N1 by deleting regions of RBM10 corresponding to aa 2–4, 4–414, and 4–742, respectively, and expressed in ARL cells. Green, EGFP; red, RBM10 stained with anti-S1-1/RBM10 antiserum. Some S1-1 NBs are indicated by arrows. Scale bar: 10 μm.

The location of the tentative NBTS was first examined in the region of aa 411–680 using deletion constructs 1–4 (Figure 2A). The product of construct 3, having a deletion of aa 524–598, localized to S1-1 NBs, but weakly, due to the presence of another NBTS outside the region of aa 411-680 (Figure 4). In contrast, products of constructs 1, 2, and 4 having deletions at aa 411–484, 476–542, and 587–680, respectively, showed distinct localization to S1-1 NBs, indicating that the regions deleted in constructs 1, 2, and 4 are not involved in NB-localization; i.e., these regions do not contain NBTSs. Figure 2C (a) shows the deleted regions in the four constructs. By excluding the regions that overlap with those in constructs 2 and 4, the NBTS expected from construct 3 is concluded to reside in aa 543–586.

**Figure 2.**
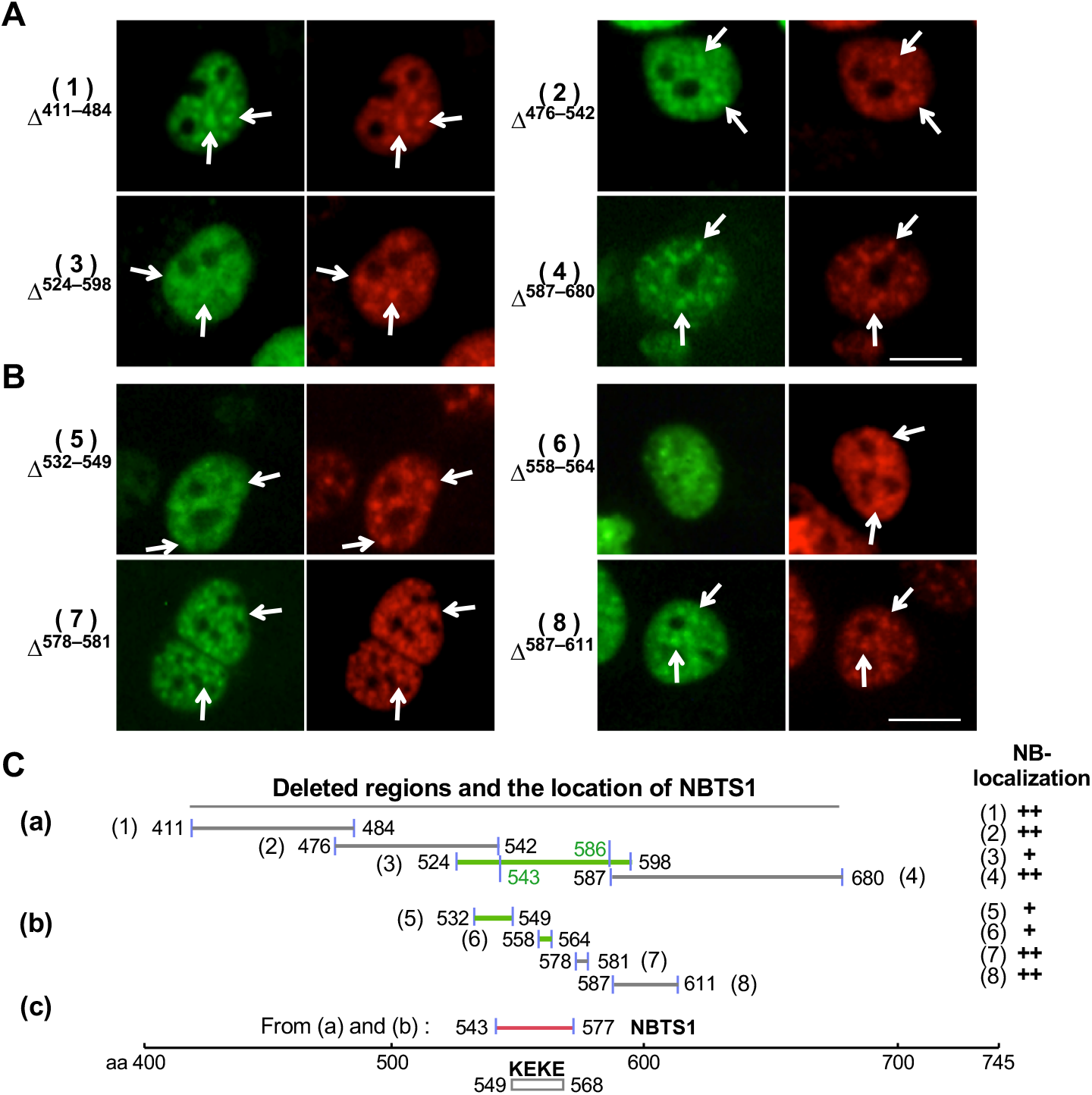
Identification of NBTS1 in aa 543–577. **A.** and **B.** Constructs (1) – (4) in A and (5) – (8) in B were derived from pRBM10/EGFP-N1 by deleting the regions indicated, and expressed in ARL cells. Green, EGFP; red, RBM10 stained with anti-S1-1/RBM10 antiserum. Some S1-1 NBs are indicated by arrows. Scale bar: 10 μm. **C.** Deleted regions and summary of results. Horizontal lines in (a) and (b) indicate deleted regions in the constructs examined in A and B, with corresponding residue numbers provided at each end. Green lines indicate constructs whose products showed weak or indistinct localization to S1-1 NBs. (c) Location of NBTS1 in RBM10. Results in (a) and (b) were combined to locate NBTS1 on RBM10. It is present in aa 543–577 (red line), which correspond to the KEKE motif region.

The NBTS in aa 543–586 was further localized using deletion constructs 5–8 [Figure 2B and 2C (b)]. In contrast to the products of constructs 7 and 8, which localized to S1-1 NBs distinctly, products of constructs 5 and 6 showed weak or indistinct NB-localization, suggesting that the NBTS resides in aa 532–577 (including the unexamined aa 565–577) [Figure 2C (b)]. By combining results shown in Figure 2C (a) and (b), the NBTS (designated as NBTS1) was assigned to aa 543–577 [Figure 2C (c)] (the assignment is verified in Figure 5). This region contains a KEKE motif in aa 549–568, characterized by alternating positive and negative amino acids [15] [Figure 3 (1)].

**Figure 3.**
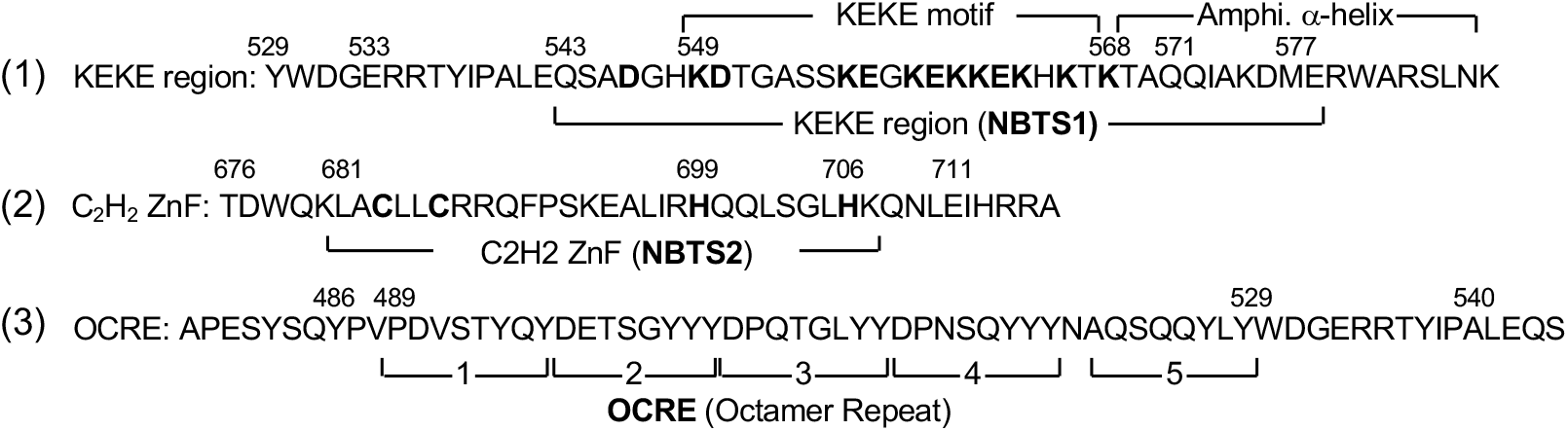
Primary sequences of the KEKE, C_2_H_2_ ZnF, and OCRE regions. (1) KEKE region. Alternating positive and negative aa in the KEKE region are indicated in boldface. “Amphi. α-helix” is a region predicted to form a perfect amphipathic α-helix (AAH) with five helical turns [1]. (2) C_2_H_2_ ZnF. Cysteine and histidine residues critical for ZnF function are highlighted in boldface. (3) OCRE (octamer repeat, alias NLS3). Brackets 1–5 represent five repeating octamers (8 aa) originally assigned based on hydrophilicity/hydrophobicity profiles [1]. The aa 486–540 region (aa 564–618 of the 930-aa isoform) forms a globular domain with an anti-parallel β-sheet fold [30].

### RBM10 contains a second NBTS

Importantly, as shown above, the product of construct 3 did not lose its NB-localization completely. Accordingly, another NBTS was thought to occur in the unexamined aa 681–742 region. This possible second NBTS was investigated using constructs pNB1/EGFP-C3 and pNB2/EGFP-C3, in which *NB1* and *NB2* encoded aa 476–645 and aa 476–760 of RBM10, respectively (Figure 4A). NB1 contains NBTS1 and a weak nuclear localization sequence (NLS3) [16]. Consequently, NB1 was detected not only in S1-1 NBs, but also in the nucleoplasm, as well as in the cytoplasm (Figure 4A). By contrast, NB2 harbors a strong NLS (NLS1) at its C-terminus [16], thus localizing predominantly to the nucleus. Moreover, like endogenous RBM10, NB2 localized distinctly to S1-1 NBs, thereby supporting the presence of another NBTS in aa 681–742 (Figure 4A).

**Figure 4.**
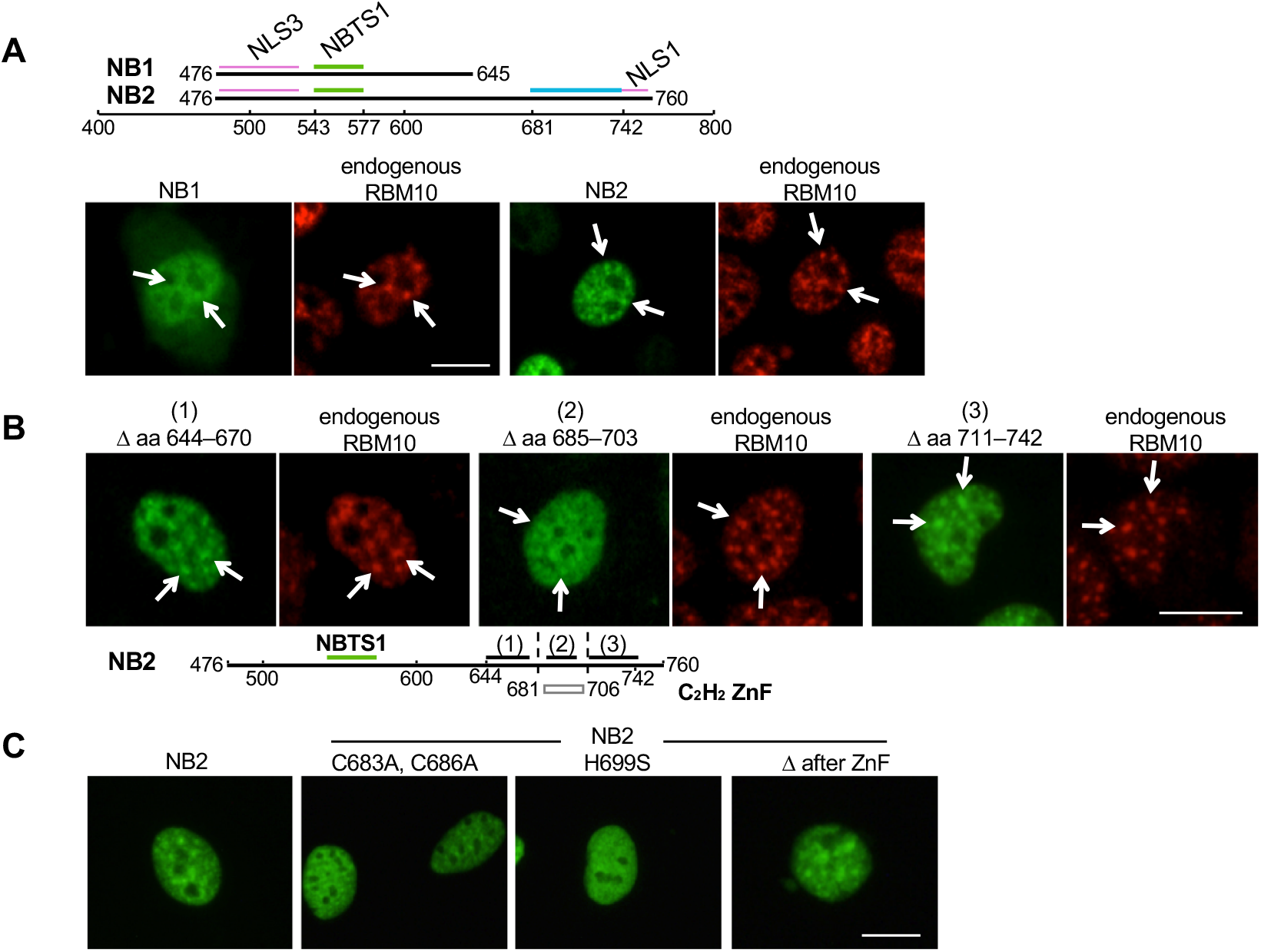
Identification of NBTS2. **A.** NB1 and NB2. RBM10 sequences encoding aa 476–645 (NB1) and 476–760 (NB2) were ligated into pEGFP-C3 and expressed in ARL cells. Green bars in the upper panel represent NBTS1 in aa 543–577 (the KEKE region). The blue bar represents the region expected to contain an additional NBTS in aa 681–742. Purple bars indicate NLS3 (aa 482–540) and NLS1 (aa 743–759, the principal NLS [16]). Lower panels show expression of pNB1/EGFP-C3 and pNB2/EGFP-C3. Green, EGFP; red, endogenous RBM10 stained with anti-S1-1/RBM10 antiserum. Representative S1-1 NBs are indicated by arrows. **B.** Determination of NBTS2. The region corresponding to aa 644–670 (control) (1), aa 685–703 (C_2_H_2_ ZnF) (2), or aa 711–742 (after-ZnF) (3) was deleted from pNB2/EGFP-C3, and constructs were expressed in ARL cells. Green, EGFP; red, RBM10 stained with anti-S1-1/RBM10 antiserum. The lower diagram indicates the location of NBTS1 (green bar), deleted regions (1) – (3) (black bars), and the location of the C_2_H_2_ ZnF (box, aa 681–706). **C.** Confirmation of NBTS2. Two cysteines at 683 and 686 were substituted with alanine or histidine at 699 was substituted with serine in the C_2_H_2_ ZnF of NB2/EGFP-C3. Green, EGFP. Scale bar: 10 μm.

Residues 681–742 comprise a C_2_H_2_-type Zn finger (ZnF) at aa 681–706 [Figure 3 (2)] and its downstream sequence (designated here as ‘after-ZnF’) at aa 707–742. These regions and an upstream control region (aa 644–670) were explored by a similar deletion analysis employing pNB2/EGFP-C3. The results were clear: only the ZnF deletion mutant showed reduced localization to S1-1 NBs [Figure 4B (2)]. It is noteworthy that due to the presence of NBTS1, the ZnF mutant still localized weakly to S1-1 NBs. To confirm the ZnF NB-targeting activity, cysteines 683 and 686, or histidine 699, which are critical for ZnF function, were replaced with alanine or serine, respectively (Figure 4C). Localizations of these two ZnF mutants to S1-1 NBs were clearly no more prominent than those of parental NB2 and the deletion mutant of after-ZnFs (Δ after-ZnF). The second NBTS in the ZnF region was designated as NBTS2 [Figure 3 (2)].

### Verification of NBTS1 and NBTS2

As shown above, the NB-targeting activity of RBM10 or NB2 was still retained after single deletion of NBTS1 (KEKE region) or NBTS2 (C_2_H_2_ ZnF), even though NB-localization was diminished by each deletion (Figures 2 and 4). Importantly, after simultaneous deletion of both NBTSs, NB-localization was no longer observed, demonstrating their synergism in NB-localization [Figure 5A (1), see also Supplementary Figure S2].

**Figure 5.**
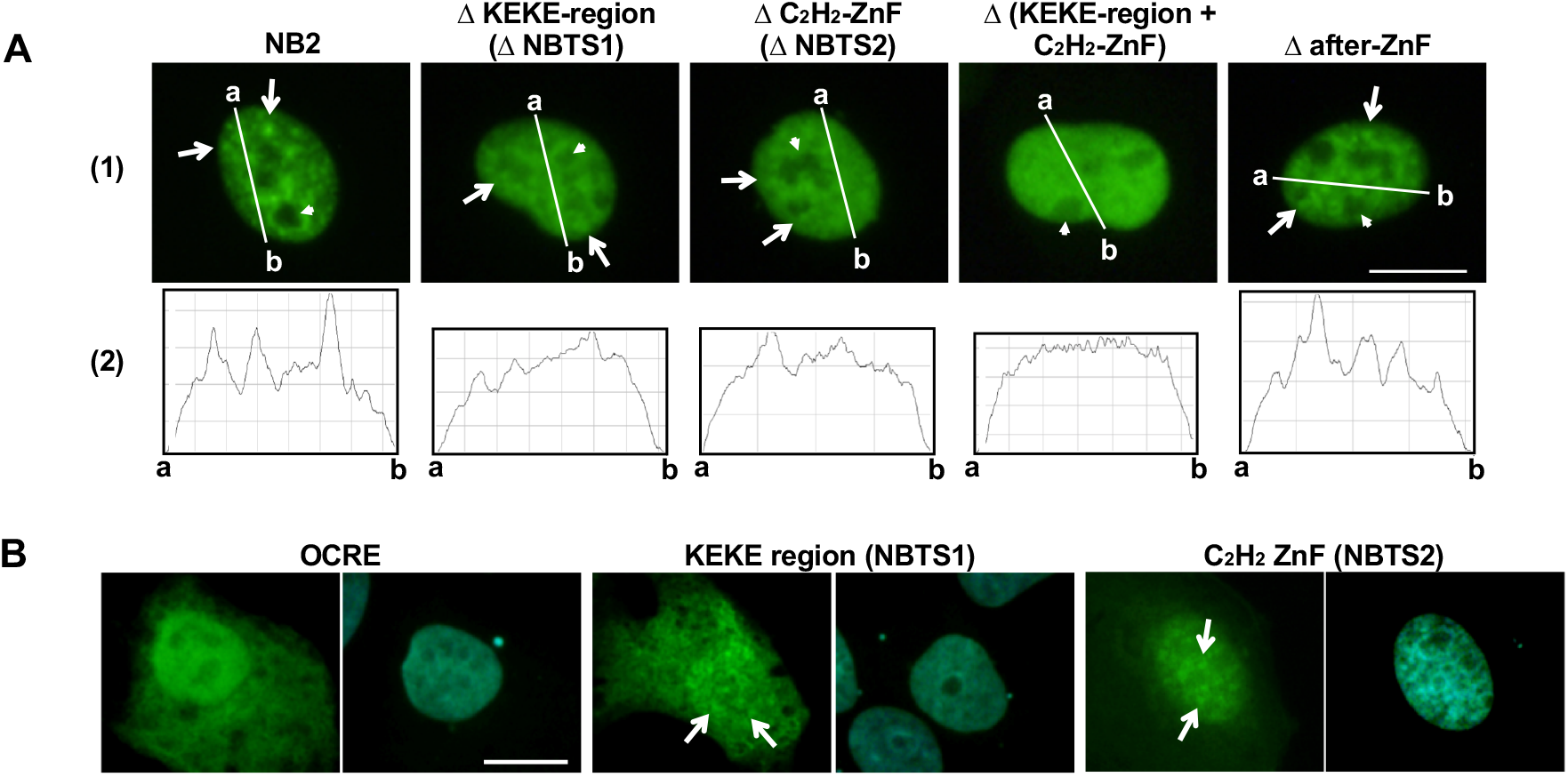
Verification of NBTSs. **A.** NB-localization of NB2 and its mutants. (1) The KEKE region (aa 524–598), the C_2_H_2_ ZnF region (aa 685–703), both the KEKE and C_2_H_2_ ZnF regions, or the after-ZnF region (aa 711–742) was deleted from pNB2/EGFP-C3, and constructs were expressed in ARL cells. Green: EGFP. Some S1-1 NBs and nucleoli are indicated by arrows and arrowheads, respectively. (2) Analysis by ImageJ Plot Profile. Nuclear distribution of NB2 and its mutants in the images in (1) was analyzed along indicated lines, drawn to traverse extra-nucleolar nucleoplasm. **B.** Verification of NB-targeting activity of the KEKE region (NBTS1) and C_2_H_2_ ZnF (NBTS2). Sequences encoding the KEKE region (aa 529–590), the C_2_H_2_ ZnF (aa 676–711), and the control OCRE region (aa 475–533) were PCR-amplified, ligated to the FLAG-tag in p3xFLAG-Myc-CMV-24, and expressed in ARL cells. Cells were stained with an antibody against FLAG (green), and nuclei were counterstained with 4’,6-diamidino-2-phenylindole (blue). Scale bar: 10 μm.

Using ImageJ Plot Profile, NB-localization was further analyzed. Since RBM10 does not localize to the nucleolus [14], analysis was performed using lines that traversed the extra-nucleolar regions [Figure 5A (2)]. In contrast with NB2 and Δafter-ZnF, NB2 mutants lacking the KEKE region (NBTS1) or the C_2_H_2_ ZnF region (NBTS2) showed blunt peaks, and the double-deletion mutant showed no distinct peaks. These results confirm the NB-targeting activities of these regions, as well as their synergism in the NB-localization.

The NBTSs were further verified by employing clones that expressed the KEKE or C_2_H_2_ ZnF region linked to the FLAG sequence. In contrast with the control octamer repeat sequence (OCRE, Figure 3(3)), the KEKE and C_2_H_2_ ZnF regions unmistakably showed NB-localization activity (Figure 5B). Based on these results, we conclude that both the KEKE region (NBTS1) and C_2_H_2_ ZnF (NBTS2) possess NB-targeting activity.

### C_2_H_2_ ZnF is essential for the AS function of RBM10

RBM10 regulates AS. We examined whether the identified NBTSs are involved in this function of RBM10. Wild-type RBM10 and RBM10 harboring mutations in NBTS1 (KEKE region) or NBTS2 (C_2_H_2_ ZnF) were expressed in RBM10-knockout HepG2 cells (Figure 6A), and their AS activities in *fas* pre-mRNA splicing were examined (Figure 6B). In this system, wild-type RBM10 promotes skipping of exon 6 in *fas* pre-mRNA [5] [Figure 6B (1)]. Exon-6 skipping of *fas* pre-mRNA was not affected by mutation of a critical KEKKE sequence in the KEKE region/NBTS1 to AAKAA, but was almost completely abolished by mutation of the CLLC sequence in the C_2_H_2_ ZnF/NBTS2 to ALLA [Figure 6B (2), see also Figure 3], demonstrating that C_2_H_2_ ZnF is essential for AS regulation by RBM10. These results indicate that the C_2_H_2_ ZnF functions both in regulation of AS and localization of RBM10 to S1-1 NBs.

**Figure 6.**
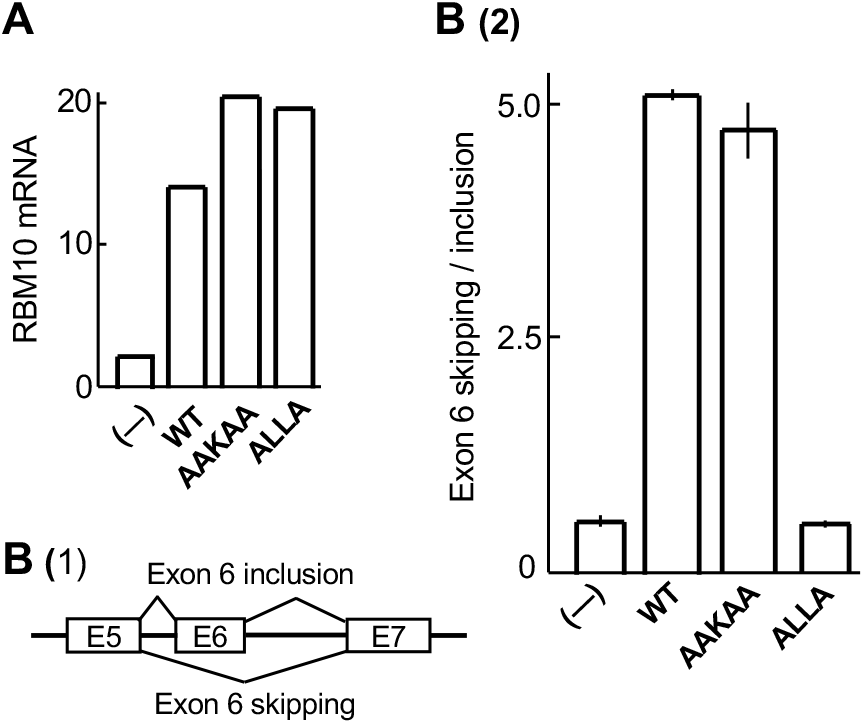
NBTS2/C_2_H_2_ ZnF is essential for regulation of AS by RBM10. RBM10-knockout HepG2 cells were transduced with lentivirus carrying wild-type RBM10 or RBM10 in which the KEKKE sequence in NBTS1 was mutated to AAKAA or the CLLC sequence in NBTS2 was mutated to ALLA. After 3 days, expression levels of mRNAs and *fas* transcripts were examined. **A.** mRNA expression levels of RBM10 and its mutants. (−), WT, AAKAA, and ALLA indicate the RBM10-knockout HepG2 cells and those transduced with wild-type RBM10 or RBM10 with the mutation of AAKAA or ALLA, respectively. **B.** (1) AS of *fas* pre-mRNA: skipping and inclusion of exon 6. (2) Skipping/inclusion ratios of exon 6. The results of two separate experiments were averaged, with deviations indicated by vertical lines at the tops of the bars. Two independent experiments in B yielded similar results, as seen in the small deviations of the averaged exon 6-skipping/inclusion values. However, RBM10 mRNA expression levels in one experiment were roughly 50% lower than those in A (data not shown), suggesting that the expressed construct proteins in both experiments were at saturation levels in performing the AS of *fas* pre-mRNA.

### RBM10 does not participate in structural organization of S1-1 NBs

We next asked whether RBM10 participates in the formation of S1-1 NBs. We previously showed that Cy3-labeled *ftz* model pre-mRNA introduced into HeLa cell nuclei by microinjection is stored in nuclear domains called TIDRs (transcription-inactivation-dependent RNA domains) when cells are incubated with actinomycin D (Act D) [17]. We further showed that TIDRs and S1-1 NBs are the same structural entities [14]. Using this experimental system, we examined whether S1-1 NBs/TIDRs form in the absence of RBM10. For this purpose, RBM10 was knocked down using specific siRNAs in HeLa cells (Figure 7A). Then, Cy3-labeled *ftz* pre-mRNA was introduced into cell nuclei, and the cells were incubated with Act D. Figure 7B demonstrates that S1-1 NBs/TIDRs formed even in the absence of RBM10, as visualized by the Cy3 fluorescence of *ftz* RNA. Thus, we concluded that RBM10 does not participate in structural organization of S1-1 NBs.

**Figure 7.**
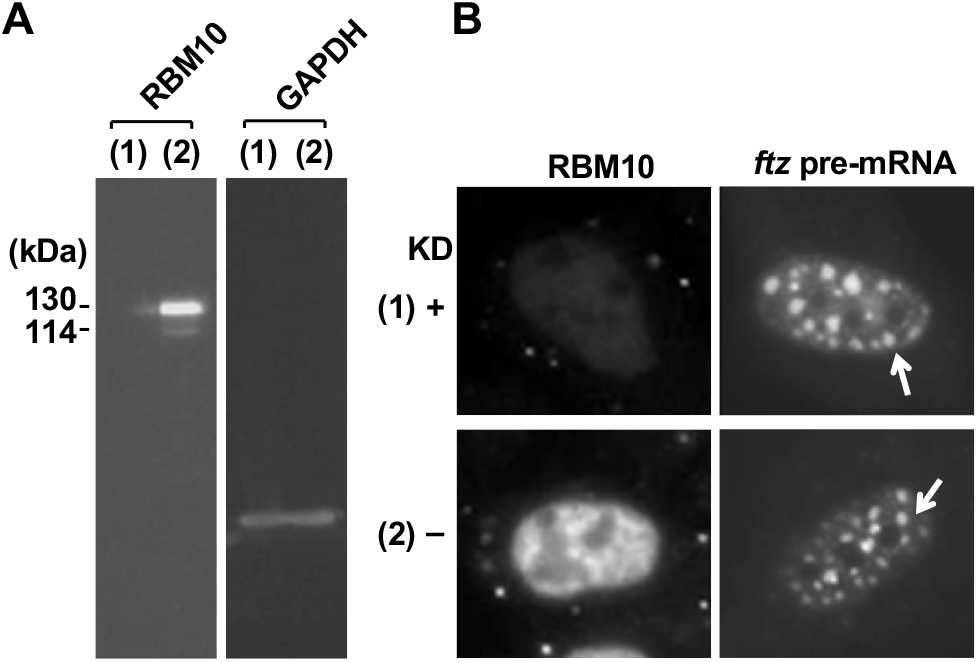
RBM10 is not essential for structural organization of S1-1 NBs. **A.** HeLa cells were treated with RBM10-specific siRNA (1) or control AllStar siRNA (2) for 3 days, and expression of RBM10 and GAPDH in these cells was examined by immunoblotting. Molecular sizes of the 930- and 852-aa isoforms are indicated in kDa on the left. **B.** Nuclei in siRNA-treated HeLa cells were microinjected with Cy3-labeled *ftz* pre-mRNA, and the cells were incubated with 5 μg/mL Act D for 2 h. Fluorescence images show RBM10 stained with anti-S1-1/RBM10 antiserum (left panels) and Cy3-labeled *ftz* pre-mRNA (right panels). Arrows indicate representative enlarged S1-1 NBs.

### RBM10 is sequestered in S1-1 NBs when transcription declines

We previously showed that localization of RBM10 to S1-1 NBs is closely associated with reduced transcription [14]. When cells are incubated with Act D, localization of RBM10 to S1-1 NBs increases in a time-dependent manner. These NBs become increasingly larger in size and fewer in number [Figure 8A (1)], while cellular levels of RBM10 remain unchanged during incubation with Act D [14]. Enlarged S1-1 NBs also form as a result of heat shock, serum starvation, or confluent cell culture, in which cellular transcriptional activity commonly decreases, and by treatment with the RNA polymerase II (pol II) inhibitor, α-amanitin [14]. In this study, we verified these previous results using other pol II inhibitors, flavopiridol and 5,6-dichloro-1-β-D-ribo-furanosylbenzimidazole (DRB). These reagents inhibit CDK9, the kinase subunit of the positive transcription elongation factor, P-TEFb, which is essential for productive transcription [18]. These two inhibitors were employed because they also act on different targets: flavopiridol inhibits cell cycle kinases, CDK1 and CDK4, and DRB inhibits casein kinase II (CK2) [19, 20]. Figure 8A (2) shows that RBM10 accumulates in S1-1 NBs upon treatment with either inhibitor, thereby suggesting that this accumulation is not due to inhibition of CDK1 and 4 or CK2, but to inhibition of P-TEFb. In support of this, DRB effects were not reproduced by the potent CK2 inhibitor, LY294002 [21] [Figure 8A (2)], or by knockdown of CK2 with specific siRNAs (data not shown). Thus, formation of enlarged S1-1 NBs with flavopiridol and DRB is due to inhibition of P-TEFb, confirming the principle that transcriptional decline induces NB-accumulation of RBM10.

**Figure 8.**
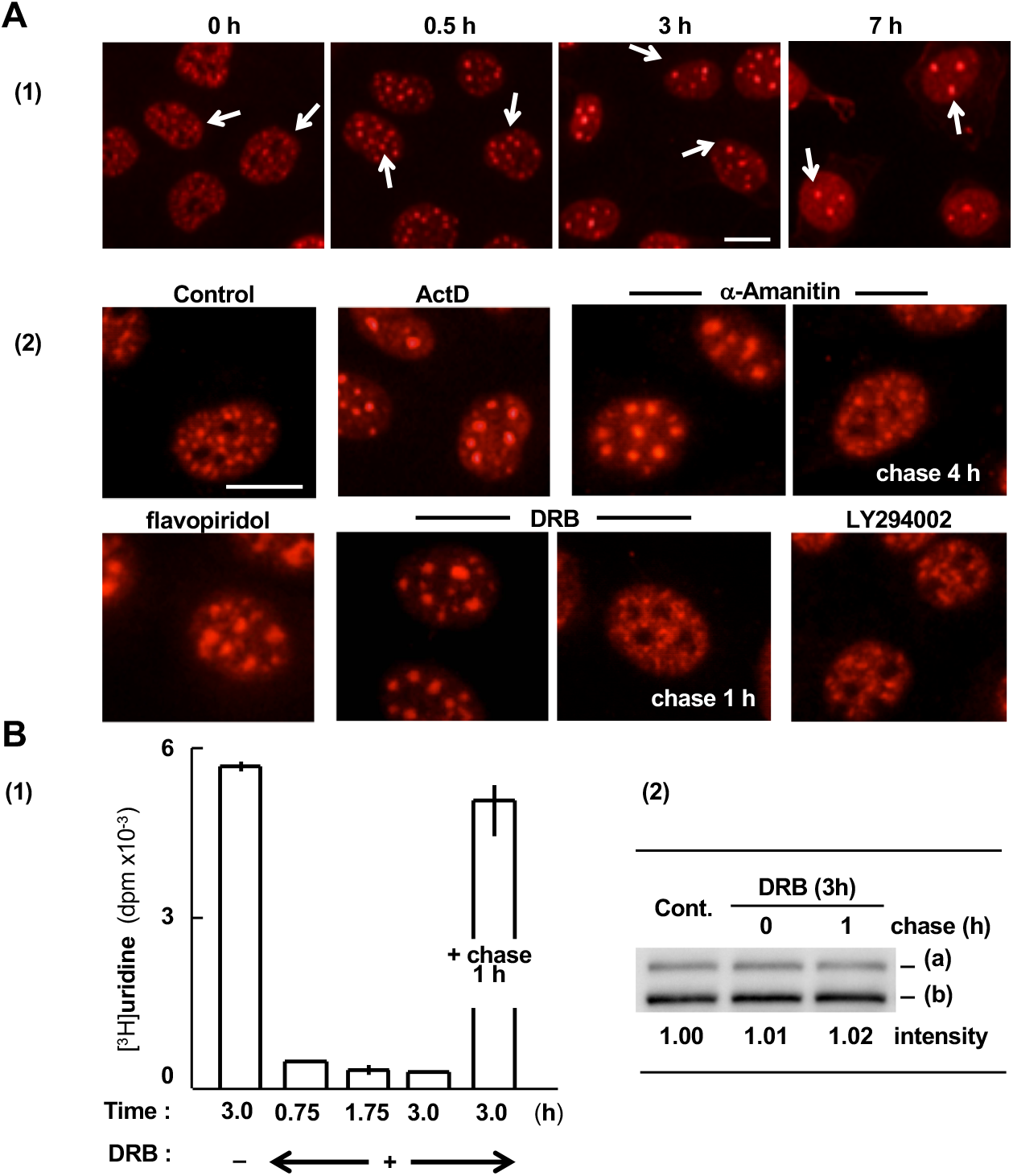
RBM10 is transiently sequestered in S1-1 NBs when transcription is suppressed. **A.** (1) Effects of Act D on S1-1 NBs. ARL cells were incubated with Act D (5.0 μg/mL) for the indicated durations and stained with anti-S1-1/RBM10 antiserum. Arrows indicate some S1-1 NBs. (2) Formation of enlarged S1-1 NBs upon suppression of transcription with various inhibitors. ARL cells were incubated with Act D, α-amanitin, flavopiridol, DRB, or LY294002 for 3 h. Note that enlarged S1-1 NBs in cells treated with α-amanitin or DRB revert to their initial state upon further incubation (chasing) in fresh media for 4 and 1 h, respectively. Cells were stained with anti-S1-1/RBM10 antiserum. Bar: 10 μm. **B.** (1) Changes in transcriptional activity. ARL cells were incubated with DRB for the indicated times, or for an additional 1 h in fresh medium without DRB (+ chase, 1h). (2) Western blotting of RBM10. ARL cells were treated with DRB as in (1). Band intensity of RBM10 and its 930- and 852-aa isoforms (a and b, respectively) are indicated.

Changes in S1-1 NBs were reversible. When cells were washed with fresh medium, the enlarged S1-1 NBs that formed in the presence of α-amanitin or DRB returned to their initial state within 4 and 1 h, respectively [Figure 8A (2)]. The difference in the length of time to reverse the effect is likely due to differences in the affinities of these inhibitors for their binding sites or because α-amanitin also triggers pol II degradation [22].

Figure 8B (1) shows the effect of DRB on cellular transcriptional activity in ARL cells. Activity decreased almost to baseline level upon treatment with DRB for 45 min. In contrast, protein levels of RBM10 remained constant in the presence of DRB for the 3-h period of observation [Figure 8B (2)], as demonstrated in 7-h treatment with Act D [14]. In addition, following a chase with fresh medium, transcriptional activity returned to nearly initial levels after 1 h, and enlarged S1-1 NBs returned to their original state [Figures A(2) DRB and 8B (1)]. Importantly, the RBM10 protein level remained constant during the chase period, too [Figure 8B (2)]. Thus, localization of RBM10 to S1-1 NBs changes in a dynamic and reversible manner according to the level of cellular transcriptional activity, and without a change in protein levels of RBM10. Dynamic, reversible localization of RBM10 to S1-1 NBs and RBM10’s lack of participation in S1-1 NB formation (Figure 7) indicate that RBM10 is a transient component of S1-1 NBs and is sequestered in these structures when cellular transcription declines.

## Discussion

The present study demonstrates that RBM10 has two NBTSs, termed NBTS1 and NBTS2, which are located in the KEKE motif and the C_2_H_2_ ZnF region, respectively. Each NBTS alone can target and localize RBM10 to S1-1 NBs; however, both NBTSs are required for increased localization of RBM10 to S1-1 NBs. Thus, NBTS1 and NBTS2 act in a synergistic manner. In RBM10, NBTS1 and NBTS2 are separated by approximately 110 amino acids and have no sequence homology (Figures 1 and 3). Thus, these NBTSs likely interact with spatially close, but distinct structures in S1-1 NBs, the geometry of which seems to form the basis for their synergistic action.

Under reduced transcription, RBM10 localizes to S1-1 NBs and these structures become enlarged. When transcriptional activity is restored, RBM10 is released from these structures and the enlarged S1-1 NBs return to their initial state ([14] and Figure 8). The present study further showed that RBM10 does not participate in S1-1 NB formation and that cellular levels of RBM10 are maintained during inhibition of transcription and during the subsequent chase period. These findings indicate that RBM10 is a transient component of S1-1 NBs and is localized to these structures when transcription decreases.

The identified NBTSs are sufficient to localize RBM10 to S1-1 NBs. All known NBs, including S1-1 NBs [14], are membrane-less compartments (for a review, see [12]); therefore, we suggest that localization of transient NB proteins to NBs occurs through association of specific sequences of the proteins (e.g., NBTSs of RBM10) with acceptor sites in corresponding NBs. This may be obvious, but it is worth noting. The dynamic, reversible NB-localization of RBM10 also suggests that RBM10 is in equilibrium between the nucleoplasm and S1-1 NBs, where the equilibrium shifts according to nucleoplasmic concentrations of pre-mRNAs.

The C_2_H_2_ ZnF/NBTS2 is bi-functional. It is not only important for NB-targeting and localization, but is also essential for regulation of pre-mRNA AS by RBM10. Hernández et al. reported that deletions from the C-terminus, including the G-patch and C_2_H_2_ ZnF, result in progressive loss of AS regulatory activity of RBM10, indicating that RBM10 integrity is required for its full function [23]. Our results support their study and further demonstrate that substitution of the two critical cysteine residues of the C_2_H_2_ ZnF with alanine is sufficient to completely abolish the AS regulatory activity of RBM10.

How the ZnF performs the dual functions of NB-targeting and AS regulation is an intriguing question. When cellular transcription declines, RBM10 decreases its AS regulatory activity due to less abundant nucleoplasmic substrate pre-mRNAs. It is presumed that when RBM10 or its C_2_H_2_ ZnF is not engaged in AS due to the absence of bound pre-mRNA, the NB-targeting activity of C_2_H_2_ ZnF acts to localize RBM10 to S1-1 NBs. Accordingly, the C_2_H_2_ ZnF is thought to act as a sensor of substrate pre-mRNA levels in the nucleoplasm, thereby determining which function, AS regulation or NB-targeting, it serves.

The *RBM10* gene resides on the X chromosome, and one of its copies is inactivated by X-inactivation in females [24. 25, 26]. This implies that a double concentration of RBM10 is unfavorable to cells and that the cellular level of active RBM10 must be controlled. In this regard, sequestration of RBM10 in S1-1 NBs may be a mechanism to rapidly, efficiently and reversibly regulate active nucleoplasmic levels of RBM10. When transcription decreases or RBM10 is not engaged in AS, RBM10 is sequestered from the nucleoplasm and stored in S1-1 NBs. When transcription increases, RBM10 dissociates from S1-1 NBs, and is promptly used for AS. We presume that this sequestration prevents nucleoplasmic RBM10 levels from becoming excessive, thereby avoiding possible unfavorable effects produced by an excess of RBM10.

Dysfunctions of RBM10 lead to various RBM10-related diseases [11]. Specific defects in NBTS activity may also cause diseases. Further studies are required to clarify the molecular mechanism that regulates sequestration of RBM10 in S1-1 NBs, including identification and characterization of acceptor sites for RBM10 in S1-1 NBs and the core proteins that comprise S1-1 NBs.

## Experimental Procedures

### Cells and cell culture

HeLa cells derived from human cervical carcinoma, HepG2 cells derived from human hepatocellular carcinoma, and HEK293T cells derived from human embryonic kidney and transformed with the SV40 large T-antigen were cultured in Dulbecco’s Modified Eagle Medium (Nissui). ARL cells (rat liver epithelial cells; Japan Health Sci. Found., JCRB0248) were cultured in Williams’ Medium E (Gibco BRL). Media were supplemented with 5% fetal calf serum (Gibco BRL), 100 μg/mL streptomycin, and 100 IU/mL penicillin. The concentration of fetal calf serum in the HepG2 cell culture was 10%. In inhibition experiments, ARL cells were incubated with 5 μg/mL Act D, 50 μg/mL α-amanitin, 0.5 mM DRB, or 0.2 μM flavopiridol.

### Immunostaining

Immunofluorescence analysis was performed as described previously [14]. RBM10 and S1-1 NBs were stained with anti-S1-1/RBM10 antiserum ([14]; 1:250 working dilution) and rhodamine-conjugated goat anti-rabbit IgG (1:200 dilution; ICN, 55662). Stained cells were examined using a fluorescence microscope (Olympus, type IX70; numerical aperture, 0.4; and objective lens, ×20). Data were captured using image acquisition software (DP Controller). Images were processed for contrast and/or brightness.

### Immunoblotting

Western blotting was performed as previously described [14]. Total protein (equivalent to 0.2 μg of cellular DNA) was electrophoresed on 10% sodium dodecyl sulfate-polyacrylamide gels (e-PARGEL; Atto).

### Primers, RBM10 mutant plasmids, and transfection

PCR primers used to prepare various constructs are listed in Table S1. Each primer contained a restriction sequence at one end for ligation of the amplified sequence into a corresponding vector. PCR conditions were as follows: elongation (30 cycles, 5 s/kb) at 72°C, denaturation (10 s) at 95°C or 98°C, and annealing (15 s) at 3.5–5.0°C below the lower T_m_ of each primer set. PCR products were digested with the corresponding restriction enzymes, purified, and ligated into pEGFP-N1, pEGFP-C3 (Clontech), or p3xFLAG-Myc-CMV-24 (Sigma-Aldrich). Competent JM109 *Escherichia coli* cells (Takara) were transformed with the constructs. Resulting colonies were cultured, and recombinant plasmids were isolated using the Pure Yield Plasmid Miniprep System (Promega). Sizes of purified plasmids were verified by agarose gel electrophoresis, and DNA sequences were determined with a 3130×1 DNA Sequencer (ABI). Transfection was performed using Lipofectamine LTX and PLUS Reagent (Invitrogen) according to the manufacturer’s recommendations.

### Constructs and mutagenesis of RBM10

The constructs ND1, ND11, and ND15 were produced by sequential deletion from pS1-1(RBM10)/EGFP-N1 [14], as described previously [16]. To generate pNB1/EGFP-C3 and pNB2/EGFP-C3, sequences encoding aa 476–645 and aa 476–760 of RBM10 were PCR-amplified and ligated into the *Sac*I and *Pst*I sites of pEGFP-C3. Deletion or substitution mutagenesis of these plasmids was performed using a PrimeSTAR Mutagenesis Basal Kit (Takara) and primers listed in Table S1. Sequences encoding aa 475–533, the KEKE region (aa 529–590 or aa 529–571), and the C_2_H_2_ ZnF region (aa 676–711) were similarly amplified (Table S1) and ligated into the *Hind*III and *Kpn*I sites of the p3xFLAG-Myc-CMV-24 vector. The two KEKE constructs yielded similar results (Figure 5B).

### RBM10 knockdown

HeLa cells were transfected with 40 nM specific human RBM10-siRNA5 or control AllStar siRNA (Qiagen) using Lipofectamine 2000 (Invitrogen), and incubated for 3 days, in accordance with the manufacturer’s protocol. A second specific siRNA, which was designed to target a different region of RBM10 mRNA, yielded similar results, indicating that the results were not due to an off-target effect.

### Nuclear injection of *ftz* pre-mRNA

When HeLa cells reached approximately 80% confluency, Cy3-labeled *ftz* pre-mRNA was microinjected into the cell nuclei using a Microinjector 5170 (Eppendorf), as previously described [17]. Cells were incubated for 2 h in the presence of Act D (5.0 μg/mL). Enlarged S1-1 NBs/TIDRs were observed by fluorescence microscopy [17].

### RBM10 knockout

RBM10-knockout HepG2 cells were generated using the CRISPR/Cas9-mediated genome editing system. Two guide RNAs (cgttcatatcctcgcgagta and cagccgagaccacgactacc) designed to target exon 3 (14-bp apart) were cloned into pX330-U6-Chimeric_BB-CBh-hSpCas9 (Addgene), according to the manufacturer’s instructions. HepG2 cells were transfected with these plasmids together with pEF-puromycin using Lipofectamine 3000 (Thermo Fisher Scientific Inc.) and incubated with 1 μg/mL puromycin for 2–3 days to obtain cell colonies. Knockout of RBM10 in the cloned cells was confirmed by Western blotting using anti-S1-1/RBM10 antiserum and by DNA sequencing.

### Generation of lentiviruses expressing wild-type and mutant RBM10

RBM10 mutants harboring mutations in NBTS1 (mutation of KEKKE to AAKAA) or NBTS2 (mutation of CLLC to ALLA), as well as a double mutant (to AAKAA/ALLA), were prepared by double-stranded plasmid mutagenesis using the primer pairs listed in Table S1 and a Quikchange Site-Directed Mutagenesis Kit (Agilent Tech.). After verification by DNA sequencing, each RBM10 cDNA was ligated into the *Xho*I and *Not*I sites of the lentiviral vector, pCSII-EF-MCS-IRES-Venus (a gift from Dr. Hiroyuki Miyoshi, RIKEN, Tsukuba, Japan). Lentivirus expressing wild-type or mutant RBM10 was produced by co-transfection of 293T cells with pCSII-EF-3xFLAG-wild type or -mutated RBM10, pCMV-VSV-G-RSV-Rev, and pCAG-HIVgp [27].

### Assessment of transcriptional activity

ARL cells at ~80% confluency in 3-cm plates were incubated with or without DRB (0.5 mM). [^3^H]-uridine was added to the cultures 30 min before reaction termination. Cells were lysed in 0.2 mL of Tris-HCl /0.2 M NaCl/0.1% SDS (pH 8.0). Plates were rinsed with 0.3 mL of Tris-HCl/0.2 M NaCl (pH 8.0). Combined cell lysates were mixed with three volumes of ethanol and incubated at −30°C for 30 min. Precipitates were collected by centrifugation and digested in 0.2 mL of 10 mM Tris-HCl/ 2 mM MgCl_2_ (pH 8.0), first with DNase I (10 *μ*g) and RNase A (5 *μ*g), and then with proteinase K (10 μg), each for 12 h. Aliquots of the digests were mixed with scintillation cocktail to measure radioactivity.

## Supporting information

This supplemental file will be used for the link to the file on the preprint site.

## Acknowledgments

SX and LW were recipients of Human Resource Development Scholarships from the Japan International Cooperation Center (JICE), Japan.

## Author Contributions

Conceptualization, AI; Methodology, AI, KNa, and TT; Validation, TY, KNi, NY; Investigation, LW, SX, HKu, MK, KT, HKo and ZH; Writing – Original Draft Preparation, AI; Writing – Review and Editing, AI and NY, and Supervision, TT, KNa, KS and AI.

## Supplementary data

Supplementary data are available at IJMS online.

