## Supplementary material for "Sequestration of RBM10 in Nuclear Bodies: Targeting Sequences and Biological Significance": This supplemental file will be used for the link to the file on the preprint site.

### Supporting information

#### Table S1. Primers

Melting temperatures ($T_m$s) of forward (Fw) and reverse (Rev) primers are indicated in parentheses on the right. Note that the 5′ portions (terminal 6–8 nucleotides in the deletion primers or the nucleotides shown in red in the substitution primers) do not participate in the hybridization with corresponding templates in PCR. Residue numbers are those of RBM10.

**1. Primers for deletion or substitution using the PrimeStar Mutagenesis Basal Kit** (Figures 2A and B)

The segments shown in red allow PCR products to circularize automatically for subsequent direct transformation in the PrimeStar Mutagenesis.

1-a) **Preparation of constructs 1–8** (deletion from the pS1-1(RBM10)/EGFP-N1 template) ($T_m$, °C)

1. aa411–484 Fw: 5′gtgggacacagtaccctgttcctgatgt3′; Rev: 5′ggtactgtgtcccaccctcaaggga3′ (55.5 and 58.8)
2. aa476–542 Fw: 5′ggccacccagtctgctgatggacataag3′; Rev: 5′gcagactgggtggccggtggtaagg3′ (54.8 and 58.5)
3. aa524–598 Fw: 5′atgctcagagtgctctacgagatgatg3′; Rev: 5′gagcactctgagcattgtagtaatactgg3′ (52.5 and 52.8)
4. aa587–680 Fw: 5′tcaacaagctagcctgtctgctctgc3′; Rev: 5′aggctagcttgttgagactgcgggc3′ (55.8 and 57.3)
5. aa532–549 Fw: 5′actgggatgacacagggcatcatca3′; Rev: 5′ctgtgtcatcccagtataggtactgctg3′ (55.5 and 55.5)
6. aa558–564 Fw: 5′aaaggagcataagacgaagacggcc3′; Rev: 5′gtcttatgctcctttgatgatgcccc3′ (53.8 and 53.9)
7. aa578–581 Fw: 5′aggacatgcgcagtctcaacaagcaa3′; Rev: 5’gactgcgcatgtccttggcaatctgt3′ (54.5 and 54.3)
8. aa587–611 Fw: 5′tcaacaaggcagatgcaggctatg3′; Rev: 5’catctgccttgttgagactgcgggc3′ (56.3 and 57.3)

1-b) **Primers for deletion from or substitution in the pNB2/EGFP-C3 template** (Figures 4B and C)

Δaa644–670 Fw: 5′gaccgccgggaagaaaagctcacag3′; Rev: 5′ttcccggcggtcatcactggcc3′ (54.8 and 56.2)

Δaa685–703 Fw: 5′ctgtctgggactccacaagcaaaacct3′; Rev: 5′ggagtcccagacaggctagcttctgcc3′ (56.6 and 57.6)

Δaa711–742 Fw: 5′caaaaccttcgcagagagaagtatggtatcc3′; Rev: 5′ctctgcgaaggttttgcttgtggagtc3′ (56.1 and 54.3)

C683A, C686A Fw: 5′cgcactgctcgcacgccgccagtttcccag3′; Rev: 5′cgtgcgagcagtgcggctagcttctgccagtctg3′ (59.8 and 59.3)

H699S Fw: 5′atccggtcacagcagctctctggactcc3′; Rev: 5′ctgctgtgaccggatgagcgcctcct3′ (58.3 and 60.2)

**2. Primers for construction of pNB1/EGFP-C3 and pNB2/EGFP-C3** (Figure 4A)

The tags indicated in red contain a *Sac*I or *Pst*I restriction site. Amplified sequences encoding aa 476–645 (NB1) or 476–760 (NB2) of RBM10 were ligated between the *Sac*I and *Pst*I sites of pEGFP-C3.

| | | |
|---|---|---|
| NB1 & NB2 Fw: | 5′tctcgagctcagtcccactgccccagagtc3′ | (62.6) |
| NB1 Rev: | 5′tcagctgcaggctagggcggtcatcactgg3′ | (61.5) |
| NB2 Rev: | 5′tcagctgcaggtacttcctccgtttgggctctg 3′ | (61.9) |

**3. Primers for construction of p3×FLAG plasmids encoding aa 475–533 (OCRE), 529–571 or 529–590 (KEKE region), or 676–711 ($C_2H_2$ ZnF) of RBM10** (Figure 5B). The segments in red contain *Hind*III or *Kpn*I restriction site sequences. Amplified sequences were ligated between the *Hind*III and *Kpn*I sites of the p3×FLAG vector.

| | | |
|---|---|---|
| aa475–533 Fw: | 5′tgacaagcttgcgaccagtcccactgcccca3′ | (62.4) |
| aa475–533 Rev: | 5′gactggtaccgactccccatcccagtataggtactgc3′ | (62.7) |
| aa529–571 Fw: | 5′tgacaagcttgcgtactgggatggggagcggc3′ | (62.9) |
| aa529–571 Rev: | 5′gactggtaccgattgggccgtcttcgtcttatgc 3′ | (62.0) |
| aa529–590 Fw: | 5′tgacaagcttgcgtactgggatggggagcggc3′ | (62.9) |
| aa529–590 Rev: | 5′gactggtaccgagttttctttttgcttgttgagactgcgg 3′ | (63.0) |
| aa676–711 Fw: | 5′tgacaagcttgcgacagactggcagaagctagcct3′ | (62.6) |
| aa676–711 Rev: | 5′gactggtaccgactcaaggttttgcttgtggagtcca3′ | (62.6) |

**4**. **Primers for generation of lentiviruses expressing wild type and mutant RBM10** (Figure 6)

| | | |
|---|---|---|
| KEKKE to AAKAA | Fw: | 5′tgtcttggtcttgtgcttcgccgccttcgctgcgccctccttcgagggtgcc3′ |
| KEKKE to AAKAA | Rev: | 5′ggcaccctcgaaggagggcgcagcgaaggcggcgaagcacaagaccaagaca3′ |
| CLLC to ALLA | Fw: | 5′gaactggcgtcgggcgagcagagcggccagcttctgc3′ |
| CLLC to ALLA | Rev: | 5′gcagaagctggccgctctgctcgcccgacgccagttc3′ |

#### Supporting Figures

##### Figure S1 Anti-S1-1/RBM10 antiserum recognizes an epitope within aa 8–63 of RBM10.

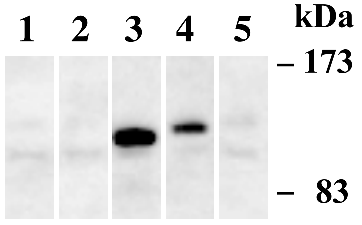


aa 1–63 (lane 1), 8–63 (lane 2), 60–136 (lane 3), or 135–169 (lane 4) was deleted from pS1-1 (aa 1–742)/EGFP-N1, which encodes aa 1–742 of RBM10 [16], and the resulting constructs were expressed in ARL cells. Using lysates of these cells, the epitope region in RBM10 recognized by the anti-S1-1/RBM10 antiserum was analyzed by immunoblotting. The antiserum was raised in a rabbit against an aa 1-166 region of the 852-aa RBM10 isoform [14]. The results indicate that the epitope recognized by the antiserum is present within aa 8–63 of RBM10. Two faint bands detected in the lysate of cells that were transfected with control pEGFP vector alone (lane 5) are attributable to the endogenous RBM10 isoforms of 852 and 930 aa residues, respectively.

**Figure S2 Enlarged images of Figure 5A (1).**

**NB2** **Δ (KEKE-region + $C_2H_2$-ZnF)**

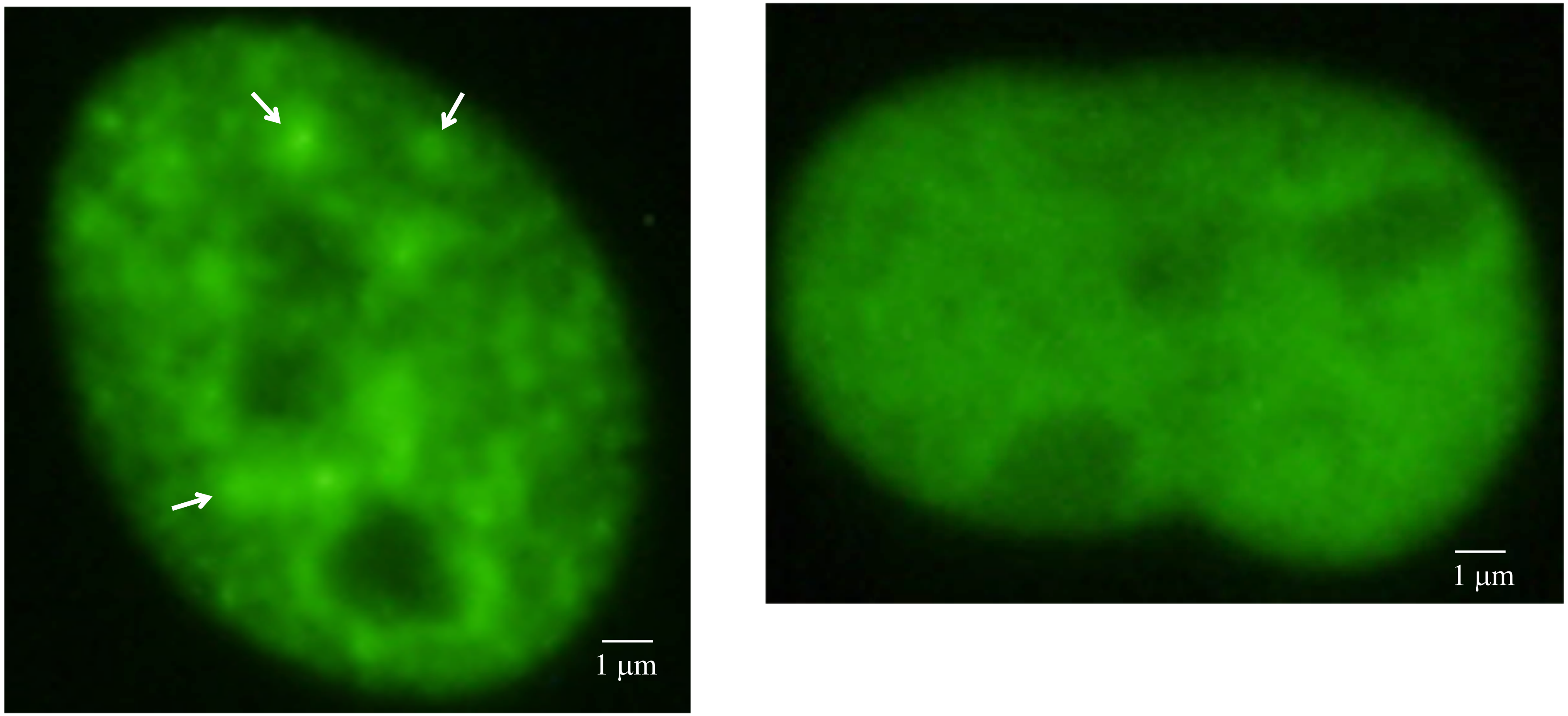


**Enlarged images of those presented in Figure 5A (1) are shown. NB2 (left) and its double deletion mutant lacking the KEKE and $C_2H_2$ ZnF regions (right) are presented. Representative S1-1 NBs are indicated by arrows.**
